# History of rare diseases and their genetic causes - a data driven approach

**DOI:** 10.1101/595819

**Authors:** Friederike Ehrhart, Egon L. Willighagen, Martina Kutmon, Max van Hoften, Nasim Bahram Sangani, Leopold G.M. Curfs, Chris T. Evelo

**Affiliations:** Department of Bioinformatics - BiGCaT, NUTRIM School of Nutrition and Translational Research in Metabolism, Maastricht University, Maastricht, The Netherlands; Governor Kremers Centre - Rett Expertise Centre, Maastricht University Medical Center, Maastricht, The Netherlands; Maastricht Centre for Systems Biology (MaCSBio), Maastricht University, Maastricht, The Netherlands

## Abstract

This dataset provides information about monogenic, rare diseases with a known genetic cause supplemented with manually extracted provenance of both the disease and the discovery of the underlying genetic cause of the disease.

We collected 4166 rare monogenic diseases according to their OMIM identifier, linked them to 3163 causative genes which are annotated with Ensembl identifiers and HGNC symbols. The PubMed identifier of the scientific publication, which for the first time describes the rare disease, and the publication which found the gene causing this disease were added using information from OMIM, Wikipedia, Google Scholar, Whonamedit, and PubMed. The data is available as a spreadsheet and as RDF in a semantic model modified from DisGeNET.

This dataset relies on publicly available data and publications with a PubMed IDs but this is to our knowledge the first time this data has been linked and made available for further study under a liberal license. Analysis of this data reveals the timeline of rare disease and causative genes discovery and links them to developments in methods and databases.

## 1. Background and summary

Descriptions of unusual diseases date back until medieval ages but rare genetic diseases are a relatively new chapter in history of medicine as genes as carriers of hereditary diseases are only discovered in the mid of the last century (see *e.g.* <u>Reflections on medicine and art</u>). Identification of the disease causing mutation in the plethora of genetic variation an individual human carries is a difficult task in the diagnosis of rare diseases. A human individual has about 25 thousand genetic variants in the exome and for the identification of the disease causing one experts cross-check with variant databases and use variant pathogenicity prediction algorithms. For the data generated in this process there are several bioinformatics workflows available to go from the raw data to the detection of the causative mutation. Within this process mapping of genetic data, identifiers and information is required in several ways. A typical workflow was described by Gilisen *et al*.^1^.

Linking information about rare diseases, their causative genes, respectively gene variants, there are many genotype-phenotype databases^2^ and some current approaches like OMIM^3^, Orphanet^4^, and DisGeNET^5^ which include provenance information - *e.g.* in form of manually curated or text mining derived literature lists. DisGeNET provides an extensive collection of linked data including a semantic model, Orphanet as well but focuses more on patient care related information. OMIM is the online version of the genetic (mendelian) disease encyclopaedia probably consulted most by clinicians. It provides information in form of a literature list with a disease or a gene and provide gene-disease mapping spreadsheets, *e.g.* morbid map. None of them provides the link of one gene to one disease including the one publication which described the disease first or found the link between gene and disease. Gene-disease associations are usually described by multiple references.

This study produced a mapping dataset which links rare, monogenic diseases to their causative genes (and *vice versa*), backed up by provenance, the publication which prove the genetic cause for a disease for the first time. The data is annotated with OMIM identifiers (disease), gene identifiers (HGNC^6^ and Ensembl^7^), and PubMed identifiers for the literature. Based on this spreadsheet we used a modified version of the DisGeNET semantic model to produce a resource description framework (RDF) file. Additionally, we produced a linkset format which can be used directly within a Cytoscape add-on CyTargetLinker^8^ (Cytoscape is a popular network analysis software^9^) for *e.g.* genetic variant or gene expression analysis linking genetic variants to genes^10^ or pathways^11^.

Based on information provided by this dataset, different interesting facts can be retrieved, *e.g.* timeline of discovery of genes responsible for rare diseases, the links of groups of diseases and epidemiology information.

## 2. Methods

### 2.1. Workflow

First, the information was collected, the suitable data was extracted and reviewed to create the dataset. Second, the dataset was created and made accessible in three different ways: spreadsheet, RDF, and linkset (for Cytoscape add-on). Third, the information from the dataset was used to retrieve information about the discovery of rare disease causing genes and, forth, applied in different analysis use cases. The workflow is shown in Figure 1.

**Figure 1:**
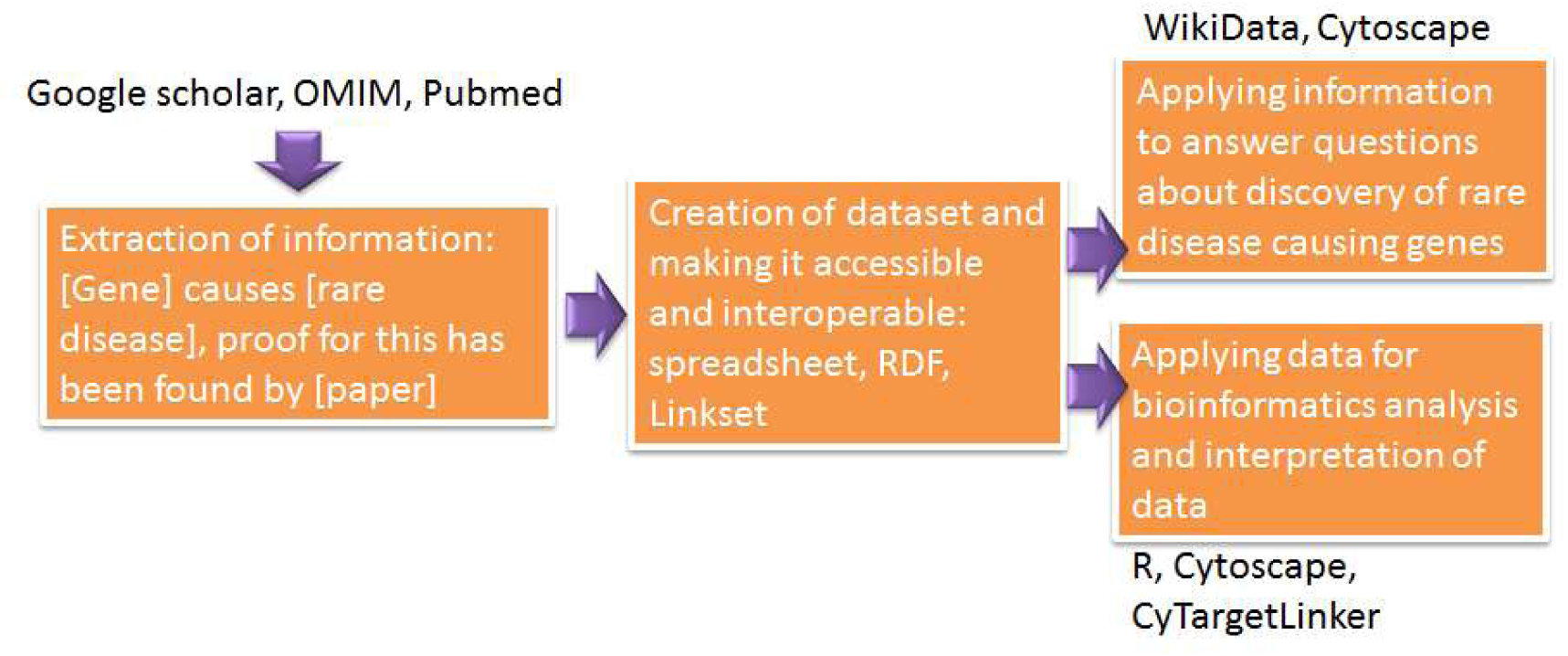
Workflow of information acquisition, creation of dataset and downstream analysis.

### 2.2. Creation of the dataset

The OMIM database was accessed for a list of all known gene-disease relationships. We manually extracted rare (about 1:1000, which is less exclusive than the EU definition^12^) monogenic diseases. The first description of the disease was manually retrieved by literature research using different resources like OMIM, PubMed, Whonamedit [https://www.whonamedit.com/], or Wikipedia. If no clear first description publication could be identified, the oldest publication in the literature list on OMIM for this disease was used. The diseases were annotated with OMIM identifiers, publications with PubMed identifiers. The causative gene for the disease was annotated with the stable gene identifier from Ensembl and the appropriate HGNC symbol. Provenance for the first publication proving that a particular disease is caused by a specific gene was manually identified based on the literature provided on OMIM and other sources (PubMed, Google Scholar). The PubMed identifier of this publication was added. Table 1 shows the structure of the final dataset.

**Table 1:** The spreadsheet data content gene-disease-provenance

| ENSID | HGNC | PMID Gene-disease | Disease OMIM ID | Disease name | PMID Disease |
| --- | --- | --- | --- | --- | --- |
| ENSG00000188536 | HBA2 | 1115799 | 604131 | Thalassemia, alpha- | 1115799 |

The dataset (version 2) contains 4565 gene-disease associations with provenance, of which 4292 are unique gene-disease associations. The difference is due to multiple PubMed identifiers for provenance as in a few cases it was not possible to identify which publication was published first. There are 3154 unique genes (according to their Ensembl identifiers) and 4166 unique diseases (OMIM identifiers) in the dataset.

### 2.3. Creation of RDF and nanopublication

To create an RDF file (according to standards as stated here: https://www.w3.org/TR/hcls-dataset/) a modified version of the DisGeNet RDF model was used. The code to create the RDF is available at: https://github.com/BiGCAT-UM/raredisease-omim/tree/master/rdf and in the attachment. The data is available in this Figshare collection: https://figshare.com/collections/Gene-Rare_Disease-Provenance_dataset_collection/4400798

The actual version which will be further discussed here is version 2: https://figshare.com/articles/Gene-RD-Provenance_V2/7718537 with the DOI: https://doi.org/10.6084/m9.figshare.7718537.v1. Nanopublications were generated from the TSV file available from Figshare using a custom Groovy script (see appendix) that makes use of nanopub-java^13^.

### 2.4. Creation of linkset for CyTargetLinker

CyTargetLinker is a Cytoscape add-on, which allows extending of networks by information given in linksets. These linksets are usually derived from external databases and contain *e.g.* gene-microRNA or drug-drug target relationships. The rare diseases (RD) linkset for CyTargetLinker was created as described in Kutmon *et al.* ^8^. Basically, we used a Java program (available here https://github.com/CyTargetLinker/linksetCreator) to convert tab delimited text files into XGMML linksets.

## 3. Data records

### 3.1. Data availability

The data (TSV) collection of the gene-RD-provenance data is available here: https://figshare.com/account/home#/collections/4400798. The currently most actual version is version 2 (DOI: <u>0.6084/m9.figshare.7718537.v1</u>). The applications of the gene-RD- provenance dataset are available on these resources: the linkset for CyTargetLinker in the <u>CyTargetLinker repository</u> and <u>RDF</u> (nanopublication). The availability of the PubMed identifiers of the publications on Wikidata^14^ was checked and if not available, a new entry was created using QuickStatements (https://tools.wmflabs.org/quickstatements/#/).

### 3.2. Software availability

The code to create the RDF from a spreadsheet or TSV is available at: https://github.com/BiGCAT-UM/raredisease-omim/tree/master/rdf. The code to create the Linkset for CyTargetLinker is available at: https://github.com/CyTargetLinker/linksetCreator. The queries to retrieve information about the publications using the pmid in Wikidata can be found here: https://github.com/egonw/pubmedWikidata or in the attachment.

## 4. Use cases and application examples

### 4.1. Bibliographic information

We queried the list of pmids in Wikidata to retrieve several different kinds of information. The SPARQL queries can be found here: https://github.com/egonw/pubmedWikidata.

#### Timeline of first descriptions of rare diseases

In this data collection the first description of a rare disease was from 1788 (Olof Ekman) about osteogenesis imperfecta. For this disease there were in the 1988 and 1989 two collagen genes identified as responsible: COL1A1^15^ and COL1A2^16^. Several first descriptions of a phenotype were later classified as separate diseases and associated with different genetic causes. In 1817 James Parkinson wrote “*An Essay on the Shaking Palsy”* describing the disease which was later named after himself. By now, there are 19 different entries in OMIM named Parkinson’s disease (or a variety of) with different genetic causes (not to be mixed up with Wolf-Parkinson syndrome which was named after Sir John Parkinson). A remarkable peak is in 1886 when Charcot-Marie-Tooth disease was described which was later sub classified in about 58 different subtypes and responsible genes. Additionally in this year first descriptions were made about Pheochromocytoma and Multiple endocrine neoplasia (each about 5 different subtypes and responsible genes). After 1901 there was no year passing without a new description of a rare disease.

#### Timeline of rare disease-gene discovery

Using the PubMed identifiers of the publication stating for the first time, that a certain disease is caused by a specific gene, Wikidata was queried (SPARQL code 10.1.6 ^17^). In Figure 2.B we plotted the number of publications per year vs timeline. In this dataset, the earliest gene-disease link is from 1967 when Seegmiller *et al.* found that a neurological disorder (which was described first three years earlier and later the name Lesch-Nyhan Syndrome was established^18^ was caused by absence of hypoxanthine-guanine phosphoribosyltransferase^19^. The discovery rate increased after invention of both Sanger sequencing and PCR, reached a plateau between about 1996 and 2006 and increased again after development of next generation sequencing. It remains to be discussed whether the decline of the discovery rates after 2013 is due to reaching the saturation (all disease causing genes found) or due to new genetic causes for syndromes may rather be allocated to previously described diseases instead of “inventing” a new disease.

**Figure 2:**
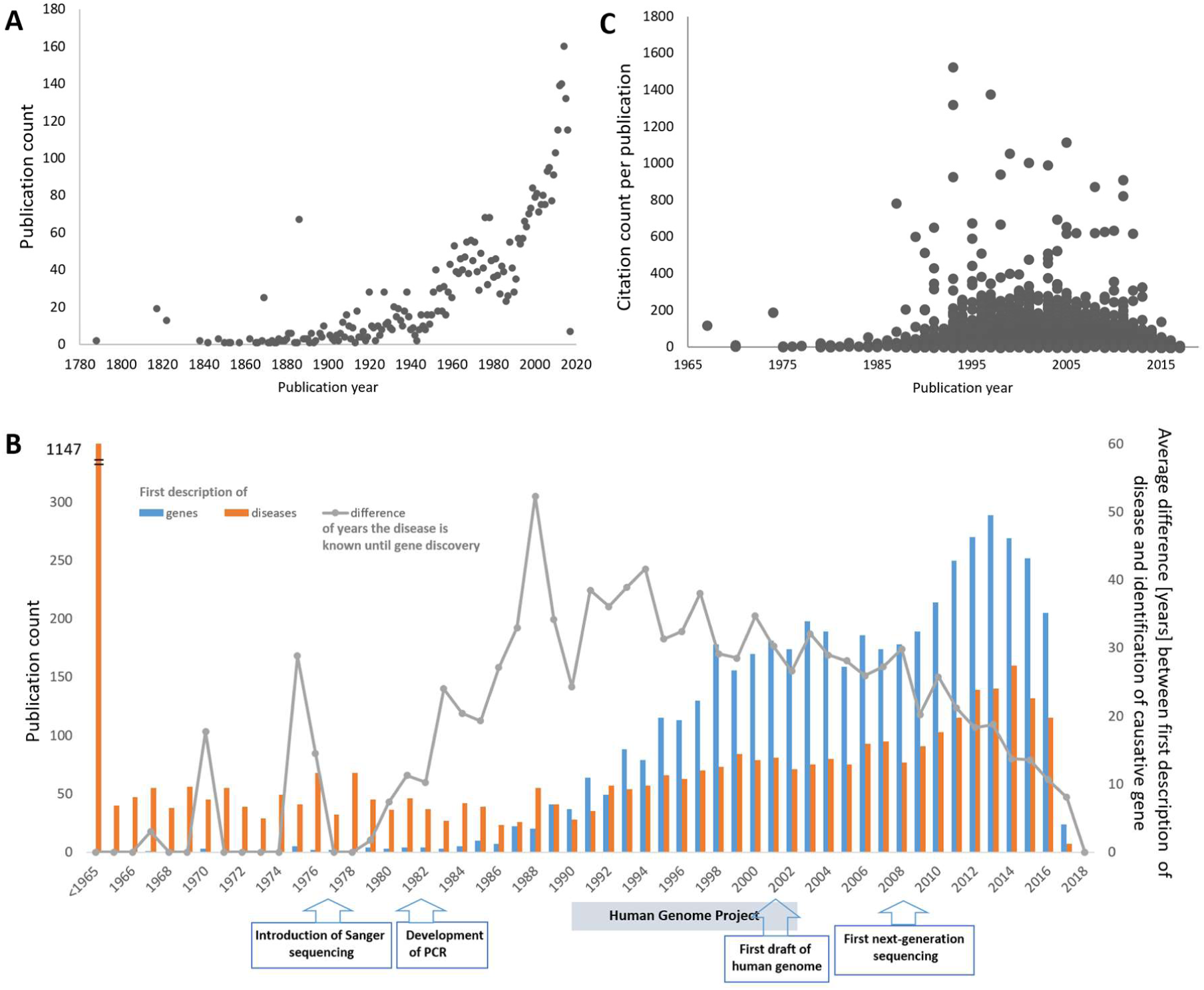
A) Number of first descriptions of rare diseases per year. B) Number of publications identifying new gene-disease links per year and the average number of years these diseases were known before the causative gene was identified. C) Citation count and year published (genes). One dot represents one publication.

In parallel, the average time since a disease was described first until the causative gene was identified was maximal in 1994 with 41.6 years and declined since then to 8 years in 2017 (Figure 2.B). After establishing of next generation sequencing in 2007, by about 2013 it become common standard that together with the description of a new genetic disease also the genetic cause is identified and published in the same document. The time span does not drop to zero due to that rare genetic causes of long known diseases (e.g. Parkinson’s or Alzheimer’s disease) are still discovered. At the moment it is speculative whether the rapidly declining peak after 2014 is due to reaching a saturation - the dataset links at the moment 3163 genes (of about 22 000 possible human genes) to one of 4166 rare genetic diseases currently known - or due to the delay of literature information reaching databases.

#### In which journals are newly discovered diseases or genes published?

The information in which journals newly discovered diseases or new causative genes for rare diseases are preferably published was retrieved from Wikidata. It was first investigated whether all these publications were available in Wikidata (https://tools.wmflabs.org/sourcemd/) and if not the paper information was uploaded (https://tools.wmflabs.org/quickstatements/#/). About 800 publications were added to Wikidata. First descriptions of rare diseases had in 3144 cases a PubMed identifier, for new genes causing a rare disease 4263 of in total 4565. To get the journals in which these papers have been published the SPARQL endpoint of Wikidata was accessed using the query given in attachment (10.2.1.). Table 2 lists the top 10 of the journals in which new diseases and new causative genes have been described. According to the data, the most important journals for publishing both, new rare diseases and newly identified genes for rare diseases are the American Journal of Human Genetics (14.9% for diseases and 26.7% for genes) and Nature Genetics (4.7% for diseases and 18.4% for genes). In total, for new rare diseases we identified 364 different journals, for genes 197. This may be due to the broader spectrum of medical disciplines and therefore journals the first observation of a now rare disease was described as well as the broader timeframe in which these were published.

**Table 2:** Journals, in which the new discovered diseases or genes causing a rare disease were published

| journal | count | % |
| --- | --- | --- |
| **First description of a new disease (100% = 3144)** |  |  |
| American Journal of Human Genetics | 457 | 14.9 |
| Nature Genetics | 145 | 4.7 |
| American Journal of Medical Genetics Part A | 137 | 4.5 |
| Journal of Medical Genetics | 136 | 4.4 |
| Human Molecular Genetics | 122 | 4.0 |
| The New England Journal of Medicine | 121 | 3.9 |
| Journal of Clinical Investigation | 78 | 2.5 |
| Neurology | 77 | 2.5 |
| The Journal of Pediatrics | 68 | 2.2 |
| Proceedings of the National Academy of Sciences of the United States of America | 59 | 1.9 |
| **First description of new genes causing a rare disease (100% = 4263)** |  |  |
| American Journal of Human Genetics | 1137 | 26.7 |
| Nature Genetics | 786 | 18.4 |
| Human Molecular Genetics | 266 | 6.2 |
| Journal of Medical Genetics | 175 | 4.1 |
| The New England Journal of Medicine | 148 | 3.5 |
| Journal of Clinical Investigation | 136 | 3.2 |
| Proceedings of the National Academy of Sciences of the United States of America | 129 | 3 |
| Science | 121 | 2.8 |
| Nature | 99 | 2.3 |
| Human Mutation | 94 | 2.2 |

#### Citation counts of the papers

The citation count of the publications was retrieved by querying Wikidata. Citation information in Wikidata is added with information from at least PubMed and CrossRef, e.g. via the WikiCite project^20^ which covers at this moment about 51% of publishers (Open Citations, https://i4oc.org/) and Wikidata has about 12.5% of all citations (status July 2018).

Figure 2.C shows the citation count distribution across the publication year for the first description of a causative gene for a rare disease. The mean citation count for such a paper is 56. Among the top 10 of most cited papers are several which identified rare, genetic causes for Alzheimer’s and Parkinson’s disease, Huntington’s disease, macular degeneration, Rett syndrome, Crohn’s disease, amyotrophic lateral sclerosis and chromosome 9p-linked frontotemporal dementia. High citation numbers indicate that there is a lot of research done for these diseases. Low citation numbers or none at all may be due to the newness of the finding (less than 5 years), disagreement among researchers, non-reproducibility of the result or no “interest” in terms of grants and research capacity investment. Among the true “neglected” diseases, diseases for which the genetic cause has been identified before 2013 but had been cited only once by now are e.g. 3-ketothiolase deficiency, glucose 6-phosphate isomerase deficiency, Leber congenital amaurosis, sarcosinemia or nonsyndromic oculocutaneous albinism. And many disorders which do not have a name yet and are described by their phenotype appearance (e.g. “Manifestations of X-linked congenital stationary night blindness in three daughters of an affected male: demonstration of homozygosity”). Apart from these, about 1000 gene-rare disease discoveries are not cited yet (according to the available data).

The most cited disease publications are Inflammatory bowel disease 25, early onset, autosomal recessive (4080 citations), D-2-hydroxyglutaric aciduria 2 (1598), Macular degeneration, age-related, 4 (1498), and Osteogenesis imperfecta, type XII (1031).

#### Authors

2641 authors of papers which first described a causative gene for a rare disease have an ORCID which is about 15% of all authors (<u>link to query</u>). Of these, 1637 are male (62.4%) and 986 female (37.6%) (18 individuals without sex/gender information). Table 3 gives the top 10 of authors with the most gene discoveries. 993 of first rare disease description papers (19.7 %) have an ORCID. The lower number may be due that the description of rare disease ranges further to the past in which ORCID was not available.

**Table 3:** Top ten of authors with most gene discoveries

| Author name | count | gender | nationality |
| --- | --- | --- | --- |
| Arnold Munnich | 58 | m | French |
| Gudrun Nürnberg | 44 | f | German |
| Peter Nürnberg | 38 | m | German |
| Thomas Meitinger | 35 | m | German |
| Nicholas Katsanis | 32 | m | British |
| Friedhelm Hildebrandt | 31 | m | USA |
| Jean-Laurent Casanova | 31 | m | French |
| Alexis Brice | 30 | m | French |
| Edgar A Otto | 19 | m | USA |
| Bruno Dallapiccola | 28 | m | Italian |

### 5.2. Application of the dataset for analysis

#### Network of genes causing diseases

Network analysis shows 2357 gene-disease causative relationships in which one gene causes one disease, res. One disease is caused by one gene (Figure 3.A upper left). Provenance can be by one or more publications depicted in one or more edges connecting the nodes. 446 triplets of two genes causing one disease, or one gene causing two diseases, are found, whereas in the majority of cases there is one gene causing two diseases. This may be explained that there are often two varieties of one disease, which were given separate identifiers in OMIM. 1226 genes and diseases are linked in more complex patterns (Figure 3.A bottom). Again, the majority, there is one gene responsible for multiple diseases or varieties of disease. The largest complex is around different, mainly mitochondrial disorders and their associated diseases. Here, a lot of overlap between the diseases and causative genes is observed, possibly due to the fact that several genes are contributing to the functionality of one complex.

**Figure 3:**
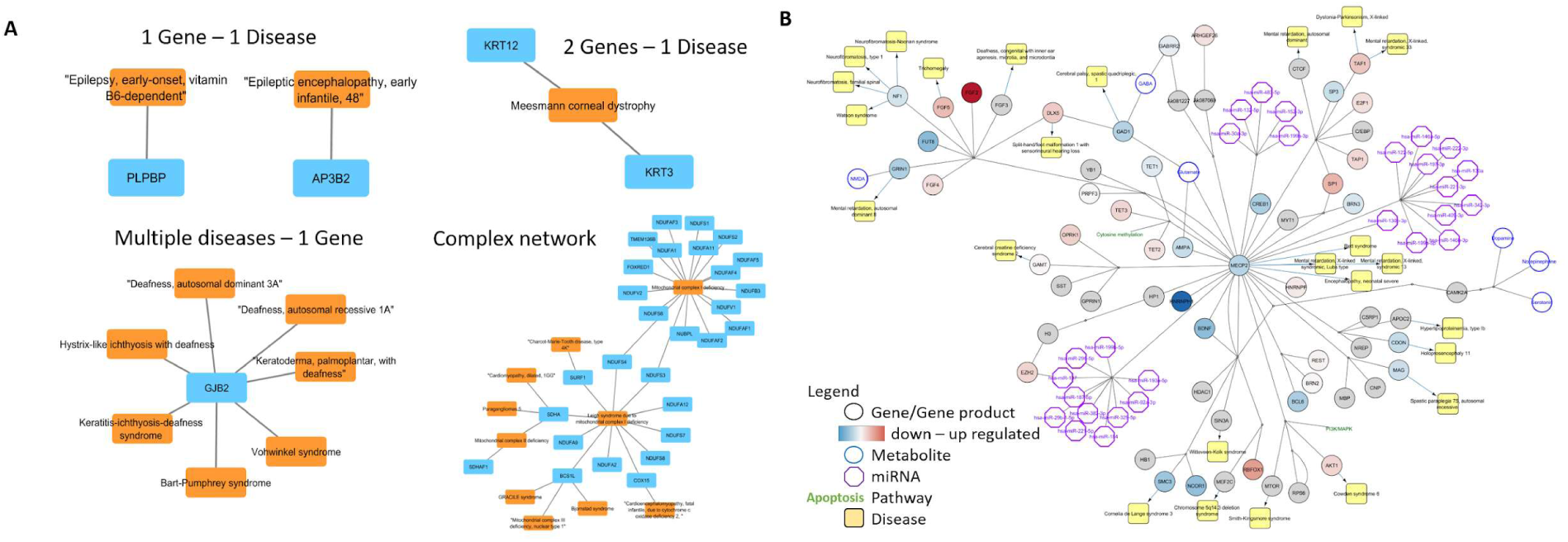
A) Network of gene-rare disease relationships. Green nodes are genes (HGNC symbols), orange nodes are diseases (OMIM disease names). B) MECP2 and associated Rett Syndrome pathway from WikiPathways, https://www.wikipathways.org/instance/WP3584 was imported as a network into Cytoscape environment using the WikiPathways app of Cytoscape. Using CyTargetLinker, the MECP2 network was then extended to predict and visualize the potential interactions between genes in the MECP2 network and other rare diseases provided by the gene-RD-Provenance_V2 linkset. The expression data was taken from Miller et al. 2019^21^ and the data was originally produced by Lin et al. 2016^22^. The network is online accessible by http://ndexbio.org/#/network/557d9e74-5209-11e9-9f06-0ac135e8bacf.

#### Integration in network analysis

Based on the information we created a linkset for the Cytoscape (popular network analysis tool) plugin CyTargetLinker (see here how to use and create these linksets https://github.com/CyTargetLinker/linksetCreator). In Figure 3.B the result of such a network extension is shown. The basic network is a rare disease pathway imported to Cytoscape from WikiPathways [https://www.wikipathways.org/instance/WP3584]. The expression data shown in the network was taken from Miller et al. 2019^21^ and the data was originally produced by Lin et al. 2016^22^. The network was then extended using the Gene-RD-Provenance_V2 linkset (available here: https://projects.bigcat.unimaas.nl/cytargetlinker/linksets/) to show which genes in the network may cause other rare genetic diseases. Examples for the identified diseases are Cornelia de Lange syndrome, Spastic paraplegia, Mental retardation, Cowden syndrome, Neurofibromatosis, or Dystonia-Parkinsonism.

### 5.3. Linking data and information with other datasets, mappings and RDF based databases

DisGeNET provides mapping datasets which allow mapping of OMIM identifiers to Concept Unique Identifiers (CUI) and Orphanet identifiers (ORPHA). Using our list of rare disease OMIM identifiers, it was possible to map 99.2% of them to a CUI and 58.0% to ORPHA.

#### DisGeNET RDF and database - identification of disease superclasses

Querying the DisGeNET database and RDF we found that 58.7% of the rare diseases are annotated with a disease superclass term (MeSH terms). The top ten of superclass terms are listed in Table 4. The majority of rare diseases are annotated with “congenital, hereditary, and neonatal diseases and abnormalities”, “nervous system disease”, and “musculoskeletal diseases”.

**Table 4:** Top 10 MeSH superclasses for the rare diseases

| Top 10 MeSH terms | Count |
| --- | --- |
| congenital, hereditary, and neonatal diseases and abnormalities | 1705 |
| nervous system disease | 945 |
| musculoskeletal diseases | 595 |
| nutritional and metabolic diseases | 549 |
| pathological conditions, signs and symptoms | 415 |
| eye diseases | 328 |
| skin and connective tissue diseases | 296 |
| cardiovascular diseases | 203 |
| hemic and lymphatic diseases | 182 |
| female genital diseases and pregnancy complications | 174 |

**Table 5:** Prevalence of the rare diseases

| Epidemiology | count |
| --- | --- |
| $>1 / 1000$ | 855 |
| 6-9 / 10 000 | 5 |
| 1-5 / 10 000 | 72 |
| 1-9 / 100 000 | 208 |
| 1-9 / 1 000 000 | 215 |
| $<1 / 1 000 000$ | 158 |
| single cases/families | 735 |
| Unknown | 354 |
| Mapping not applicable | 1963 |

#### What kind of disorders have been discovered and when?

Linking publishing date of a rare disease description with MeSH disease superclass information from DisGeNET reveals trends which class of diseases have been identified. Figure 4 shows a timeline in blocks of 20 years, in which how many rare diseases belonging to different superclasses been described.

**Figure 4:**
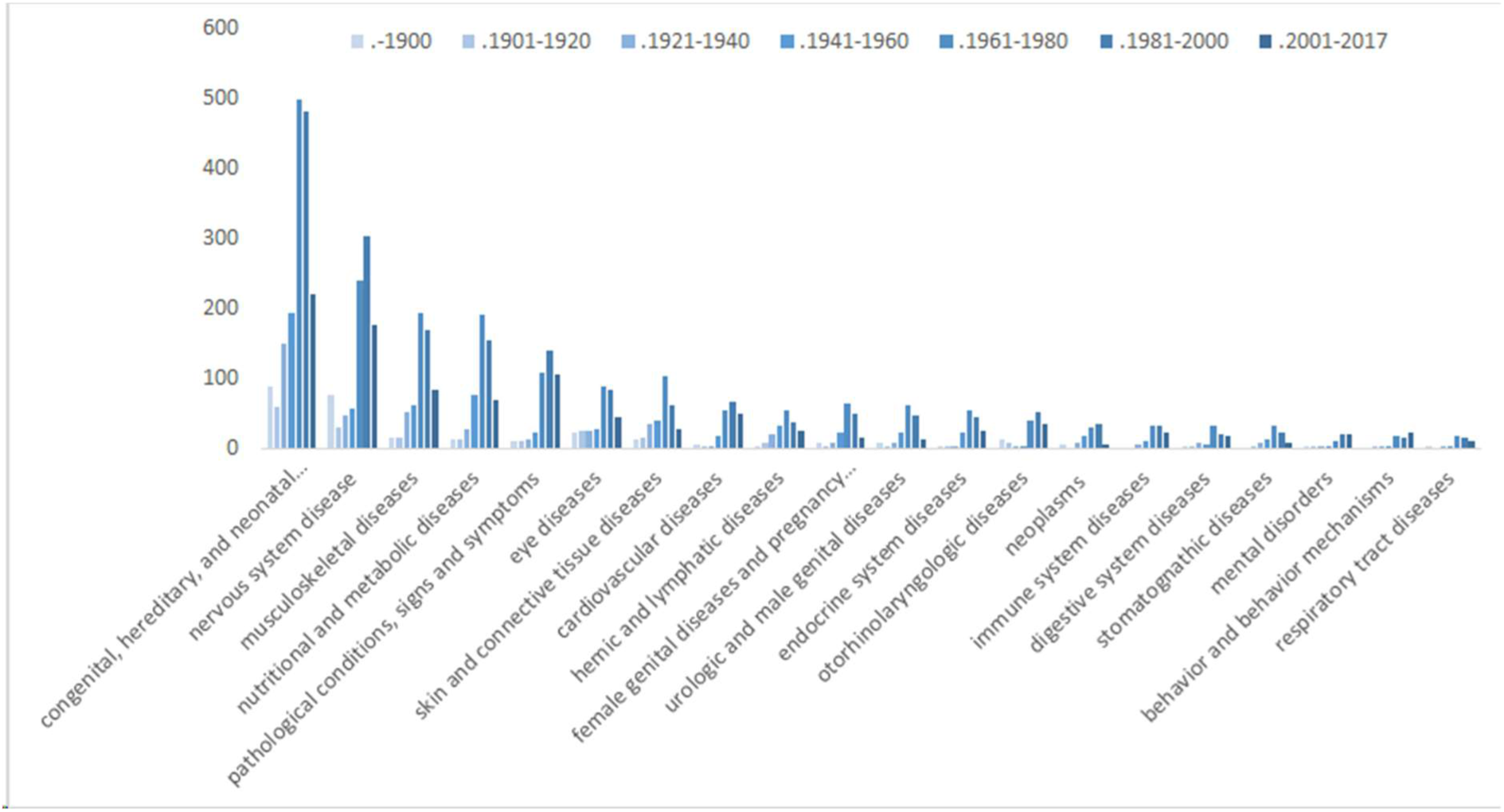
Timeline of rare disease superclass descriptions in blocks of 20 years.

#### Prevalence distribution of rare diseases

Orphanet provides data collections about disease prevalence. Linking this information via disease identifier mapping between OMIM, CUI and ORPHA shows that the majority of diseases in the list are occurring >1/1000 which would be not a rare disease. This is due to mapping between the databases and more common phenotypes (e.g. Alzheimer’s disease) caused by rare genetic mutations which we counted in as rare monogenic diseases.

## Author contributions

Friederike Ehrhart (creating dataset, data analysis, initiation, organization, application test) Chris Evelo (critical review)

Max van Hoften (creating dataset)

Leopold Curfs (critical review)

Egon Willighagen (creating RDF of dataset, R scripts)

Martina Summer-Kutmon (creating linkset for CyTargetLinker)

Nasim Bahram Sangani (Cytoscape use case)

## Acknowledgement

Thanks to creators and curators of databases and mappings.

## Supplementary material

### 10.1 R scripts

#### 10.1.1. Install dependencies

~~~
install.packages(pkgs=
  c(“googlesheets”, “plyr”, “foreach”)
)
~~~

#### 10.1.2. Create RDF in DisGeNET format

~~~
@Grapes([
  @Grab(group=‘org.nanopub’, module=‘nanopub’, version=‘1.18’),
  @Grab(‘com.xlson.groovycsv:groovycsv:1.1’)
])
import static com.xlson.groovycsv.CsvParser.parseCsv
import org.nanopub.Nanopub;
import org.nanopub.NanopubCreator;
import org.nanopub.NanopubUtils;
import org.openrdf.model.impl.ValueFactoryImpl;
import org.openrdf.model.vocabulary.RDF;
import org.openrdf.model.vocabulary.RDFS;
import org.openrdf.model.vocabulary.SKOS;
import org.openrdf.model.vocabulary.OWL;
import org.openrdf.rio.RDFFormat;
factory = ValueFactoryImpl.getInstance()
input = args[0]
counter = 0
hasSource = factory.createURI(“http://semanticscience.org/resource/SIO_000253”)
database = factory.createURI(“http://semanticscience.org/resource/SIO_000750”)
SIO000983 = factory.createURI(“http://semanticscience.org/resource/SIO_000983”)
SIO000628 = factory.createURI(“http://semanticscience.org/resource/SIO_000628”)
ncitC4873= factory.createURI(“http://ncicb.nci.nih.gov/xml/owl/EVS/Thesaurus.owl#C4873”)
DOID0050177 = factory.createURI(“http://purl.obolibrary.org/obo/DOID_0050177”)
wasDerivedFrom = factory.createURI(“http://www.w3.org/ns/prov#wasDerivedFrom”)
wasGeneratedBy = factory.createURI(“http://www.w3.org/ns/prov#wasGeneratedBy”)
Q241953 = factory.createURI(“http://www.wikidata.org/entity/Q241953”)
ECO000218 = factory.createURI(“http://purl.obolibrary.org/obo/eco.owl#ECO_000218”)
rights = factory.createURI(“http://purl.org/dc/terms/rights”)
cczero = factory.createURI(“http://creativecommons.org/publicdomain/zero/1.0/”)
p3331 = factory.createURI(“http://www.wikidata.org/prop/direct/P3331”)
figshare = factory.createURI(“https://doi.org/10.6084/m9.figshare.7718537.v1”)
def text = new File(input).getText()
def data = parseCsv(text, separator: ‘\t’)
data.each { line ->
geneID = line[“ENSID”]
hgnc = line[“HGNC”]
genePMID = line[“PMID Gene-disease”]
disease = line[“Disease name”]
diseasePMID = line[“PMID Disease”]
omimID = line[“Disease OMIM ID”]
if (geneID.isEmpty()) return
// the Disease NanoPub
counter++
nanopubIRI = “http://purl.org/nanopub/temp/np” + counter
creator = new NanopubCreator(nanopubIRI)
creator.addTimestamp(new Date())
creator.addNamespace(“np”, “http://www.nanopub.org/nschema#”)
creator.addNamespace(“orcid”, “http://orcid.org/”)
creator.addNamespace(“wdt”, “http://www.wikidata.org/prop/direct/”)
creator.addNamespace(“pav”, “http://purl.org/pav/”)
creator.addNamespace(“rdfs”, “http://www.w3.org/2000/01/rdf-schema#”)
creator.addNamespace(“dct”, “http://purl.org/dc/terms/”)
creator.addNamespace(“xsd”, “http://www.w3.org/2001/XMLSchema#”)
creator.addNamespace(“prov”, “http://www.w3.org/ns/prov#”)
creator.addNamespace(“dc”, “http://purl.org/dc/elements/1.1/”)
creator.addNamespace(“pmid”, “http://identifiers.org/pubmed/”)
creator.addNamespace(“ens”, “http://identifiers.org/ensembl/”)
creator.addNamespace(“cczero”,
“http://creativecommons.org/publicdomain/zero/1.0/”)
creator.addNamespace(“hgnc”, “http://identifiers.org/hgnc/”)
creator.addNamespace(“omim”, “http://identifiers.org/omim/”)
creator.addNamespace(“eco”, “http://purl.obolibrary.org/obo/eco.owl#”)
creator.addNamespace(“ncit”,
“http://ncicb.nci.nih.gov/xml/owl/EVS/Thesaurus.owl#“)
creator.addNamespace(“obo”, “http://purl.obolibrary.org/obo/”)
creator.addNamespace(“figshare”, “https://doi.org/10.6084/“)
omimURI = “http://identifiers.org/omim/” + omimID
omim = factory.createURI(omimURI)
creator.addAssertionStatement(omim, RDF.TYPE, ncitC4873)
creator.addAssertionStatement(omim, RDF.TYPE, DOID0050177)
creator.addAssertionStatement(omim, RDFS.LABEL, factory.createLiteral(disease))
creator.addProvenanceStatement(creator.getAssertionUri(), wasDerivedFrom,
figshare)
if (diseasePMID != null && !diseasePMID.isEmpty() && !diseasePMID.contains(“,”))
{
try {
pmidVal = Integer.parseInt(diseasePMID);
diseasePMIDURI = factory.createURI(“http://identifiers.org/pubmed/” +
diseasePMID)
creator.addProvenanceStatement(creator.getAssertionUri(), wasDerivedFrom, diseasePMIDURI)
} catch (Exception exc) {} // ignore
}
creator.addProvenanceStatement(creator.getAssertionUri(), wasGeneratedBy, ECO000218)
creator.addPubinfoStatement(rights, cczero)
egon = creator.getOrcidUri(“0000-0001-7542-0286”)
freddie = creator.getOrcidUri(“0000-0002-7770-620X”)
creator.addCreator(freddie)
creator.addCreator(egon)
trustedPub = creator.finalizeTrustyNanopub()
outputBuffer = new StringBuffer();
outputBuffer.append(NanopubUtils.writeToString(trustedPub,
RDFFormat.TRIG)).append(“\n\n”);
println outputBuffer.toString()
// the Association NanoPub
counter++
nanopubIRI = “http://purl.org/nanopub/temp/np” + counter
creator = new NanopubCreator(nanopubIRI)
creator.addTimestamp(new Date())
creator.addNamespace(“np”, “http://www.nanopub.org/nschema#”)
creator.addNamespace(“orcid”, “http://orcid.org/”)
creator.addNamespace(“wdt”, “http://www.wikidata.org/prop/direct/”)
creator.addNamespace(“pav”, “http://purl.org/pav/”)
creator.addNamespace(“rdfs”, “http://www.w3.org/2000/01/rdf-schema#”)
creator.addNamespace(“dct”, “http:purl.org/dc/terms/”)
creator.addNamespace(“xsd”, “http://www.w3.org/2001/XMLSchema#”)
creator.addNamespace(“prov”, “http://www.w3.org/ns/prov#”)
creator.addNamespace(“dc”, “http:purl.org/dc/elements/1.1/”)
creator.addNamespace(“pmid”, “http:identifiers.org/pubmed/”)
creator.addNamespace(“ens”, “http:identifiers.org/ensembl/”)
creator.addNamespace(“cczero”,
“http:creativecommons.org/publicdomain/zero/1.0/”)
creator.addNamespace(“hgnc”, “http:identifiers.org/hgnc/”)
creator.addNamespace(“omim”, “http:identifiers.org/omim/”)
creator.addNamespace(“eco”, “http:purl.obolibrary.org/obo/eco.owl#”)
creator.addNamespace(“ncit”,
“http://ncicb.nci.nih.gov/xml/owl/EVS/Thesaurus.owl#“)
creator.addNamespace(“obo”, “http:purl.obolibrary.org/obo/”)
creator.addNamespace(“figshare”, “https://doi.org/10.6084/“)
creator.addNamespace(“bc”, “http://www.bigcat.unimaas.nl/rett/”)
creator.addNamespace(“sio”, “http:semanticscience.org/resource/”)
assocURI = “http://www.bigcat.unimaas.nl/rett/as” +
Math.abs((geneID+omimID).hashCode())
assoc = factory.createURI(assocURI)
creator.addAssertionStatement(assoc, RDF.TYPE, SIO000983)
geneURI = “http:identifiers.org/ensembl/” + geneID
gene = factory.createURI(geneURI)
creator.addAssertionStatement(assoc, SIO000628, gene)
creator.addAssertionStatement(gene, RDFS.LABEL, factory.createLiteral(hgnc))
omimURI = “http:identifiers.org/omim/” + omimID
omim = factory.createURI(omimURI)
creator.addAssertionStatement(assoc, SIO000628, omim)
creator.addAssertionStatement(omim, RDFS.LABEL, factory.createLiteral(disease))
creator.addProvenanceStatement(creator.getAssertionUri(), wasDerivedFrom, figshare)
if (genePMID != null && !genePMID.isEmpty() && !genePMID.contains(“,”)) {
genePMIDURI = factory.createURI(“http:identifiers.org/pubmed/” + genePMID)
creator.addProvenanceStatement(creator.getAssertionUri(), wasDerivedFrom,
genePMIDURI)
}
creator.addProvenanceStatement(creator.getAssertionUri(), wasGeneratedBy,
ECO000218)
creator.addPubinfoStatement(rights, cczero)
egon = creator.getOrcidUri(“0000-0001-7542-0286”)
freddie = creator.getOrcidUri(“0000-0002-7770-620X”)
creator.addCreator(freddie)
creator.addCreator(egon)
trustedPub = creator.finalizeTrustyNanopub()
outputBuffer = new StringBuffer();
outputBuffer.append(NanopubUtils.writeToString(trustedPub,
RDFFormat.TRIG)).append(“\n\n”);
println outputBuffer.toString()
}
~~~

#### 10.1.3. Fetch Wikidata entity IRIs for the PubMed IDs

~~~
library(googlesheets)
library(foreach)
library(plyr)
library(rrdf)
#gs_ls(regex=“Dataset Monogenic Rare Diseases”)
omimspread <- gs_title(“Dataset Monogenic Rare Diseases - Version 2018-03-14”) omimdata = gs_read(omimspread, ws=“gene-disease-provenance”)
store = new.rdf(ontology=FALSE)
add.prefix(store, “owl”, “http://www.w3.org/2002/07/owl#”)
add.prefix(store, “pubmed”, “http:identifiers.org/pubmed/”)
add.prefix(store, “wd”, “http://www.wikidata.org/entity/”)
pubmedIDs = as.vector(unlist(omimdata[,”pubmed ID”]))
pubmedidsAsValues = ““
for(pmid in pubmedIDs) {
  pubmedidsAsValues = paste(
    pubmedidsAsValues, “ \”“, pmid, “\”\n”,
    sep=““
  )
}
template = “
PREFIX wdt: <http://www.wikidata.org/prop/direct/>
SELECT * WHERE {
  VALUES ?pmid {
${pubmedidsAsValues}
  }
  ?article wdt:P698 ?pmid
}
“
query = sub(“${pubmedidsAsValues}”, pubmedidsAsValues, template, fixed=TRUE)
print(query)
results = sparql.remote(
  “https://query.wikidata.org/bigdata/namespace/wdq/sparql“,
  query,
  jena=FALSE
)
results
adply(results, 1, function(row) {
  wkid = as.character(row[“article”])
  pmid = as.character(row[“pmid”])
  pubmedIRI = paste(“http:identifiers.org/pubmed/”, pmid, sep=““)
  add.triple(store,
    pubmedIRI, “http://www.w3.org/2002/07/owl#sameAs“, wkid
)
})
save.rdf(store, “omim_wikidata_pubmed.ttl”, format=“N3”)
~~~

#### 10.1.4. Fetch Wikidata entity IRIs for the Gene IDs

~~~
library(googlesheets)
library(foreach)
library(plyr)
library(rrdf)
#gs_ls(regex=“Dataset Monogenic Rare Diseases”)
omimspread <- gs_title(“Dataset Monogenic Rare Diseases - Version 2018-03-14”)
omimdata = gs_read(omimspread, ws=“gene-disease-provenance”)
store = new.rdf(ontology=FALSE)
add.prefix(store, “rdf”, “http://www.w3.org/1999/02/22-rdf-syntax-ns#”)
add.prefix(store, “rdfs”, “http://www.w3.org/2000/01/rdf-schema#”)
add.prefix(store, “ens”, “http:identifiers.org/ensembl/”)
add.prefix(store, “owl”, “http://www.w3.org/2002/07/owl#”)
add.prefix(store, “wd”, “http://www.wikidata.org/entity/”)
geneIDs = as.vector(unlist(omimdata[,”Gene stable ID”]))
geneidsAsValues = ““
for(id in geneIDs) {
  geneidsAsValues = paste(
    geneidsAsValues, “ \”“, id, “\”\n”,
      sep=““
  )
}
template = “
PREFIX wdt: <http://www.wikidata.org/prop/direct/>
SELECT * WHERE {
  VALUES ?geneid {
${geneidsAsValues}
  }
  ?gene wdt:P594 ?geneid}
“
query = sub(“${geneidsAsValues}”, geneidsAsValues, template, fixed=TRUE)
print(query)
results = sparql.remote(
  “https://query.wikidata.org/bigdata/namespace/wdq/sparql“,
  query,
  jena=FALSE
)
results
adply(results, 1, function(row) {
  wkid = as.character(row[“gene”])
  geneid = as.character(row[“geneid”])
  geneIRI = paste(“http:identifiers.org/ensembl/”, geneid, sep=““)
  add.triple(store,
    geneIRI, “http://www.w3.org/2002/07/owl#sameAs”, wkid
  )
})
save.rdf(store, “omim_wikidata_gene.ttl”, format=“N3”)
~~~

### 10.2. SPARQL queries

#### 10.2.1. Fetch Wikidata journalLabel for pmid

http://tinyurl.com/y86cbdyw (add whole list, export tsv)

~~~
SELECT ?journal ?journalLabel (COUNT(?pmid) AS ?count) WHERE {
  VALUES ?pmid {
“10024875”
  }
  ?article wdt:P1433 ?journal;
           wdt:P698 ?pmid.
  SERVICE wikibase:label { bd:serviceParam wikibase:language “[AUTO_LANGUAGE],en”.
}
} GROUP BY ?journal ?journalLabel
ORDER BY DESC(?count)
~~~

#### 10.2.2. Wikidata query for publication date of a paper

~~~
SELECT ?publicationDate ?pmid WHERE {
  VALUES ?pmid {
“15613”
  }
  ?article wdt:P577 ?publicationDate;
           wdt:P698 ?pmid.
}
~~~

